# ReadBouncer: Precise and Scalable Adaptive Sampling for Nanopore Sequencing

**DOI:** 10.1101/2022.02.01.478636

**Authors:** Jens-Uwe Ulrich, Ahmad Lutfi, Kilian Rutzen, Bernhard Y. Renard

## Abstract

Nanopore sequencers allow targeted sequencing of interesting nucleotide sequences by rejecting other sequences from individual pores. This feature facilitates the enrichment of low-abundant sequences by depleting overrepresented ones in-silico. Existing tools for adaptive sampling either apply signal alignment, which cannot handle human-sized reference sequences, or apply read mapping in sequence space relying on fast GPU base callers for real-time read rejection. Using nanopore long-read mapping tools is also not optimal when mapping shorter reads as usually analyzed in adaptive sampling applications. Here we present a new approach for nanopore adaptive sampling that combines fast CPU and GPU base calling with read classification based on Interleaved Bloom Filters (IBF). ReadBouncer improves the potential enrichment of low abundance sequences by its high read classification sensitivity and specificity, outperforming existing tools in the field. It robustly removes even reads belonging to large reference sequences while running on commodity hardware without graphical processing units (GPUs), making adaptive sampling accessible for in-field researchers. Readbouncer also provides a user-friendly interface and installer files for end-users without a bioinformatics background.

**Availability:** The C++ source code of ReadBouncer is available at (https://gitlab.com/dacs-hpi/readbouncer).

## Introduction

During the last decade, the invention of nanopore sequencing instruments has democratized DNA sequencing in various aspects (1, 2). For example, the small MinION devices of Oxford Nanopore Technologies provide the possibility to sequence a sample at the place of its origin, without the need to ship the sample to a laboratory (3, 4). This point-of-care sequencing ability makes nanopore sequencing attractive for applications such as pathogen detection in a clinical setting and in the field (5, 6). It also can shorten the time to detect pathogens or antimicrobial resistance (AMR) genes when using it for point-of-care testing.

While the size of the device and the easier and faster sample preparation are clear advantages, nanopore sequencing still lags the base quality of sequencing-by-synthesis instruments. However, recent improvements in base-calling algorithms showed per read accuracy exceeding 90% (7, 8). Oxford Nanopore Technologies (ONT) even claims to boost per read accuracy up to 99% with their latest R10.4 pore version (https://nanoporetech.com/accuracy).Another exciting feature of ONT’s instruments is sequencing DNA molecules in a targeted fashion. Oxford Nanopore provides an Application Programming Interface (API) that enables receiving electrical currents, measured while the molecule transverses the pore (9). These signals can be translated into sequence space and analyzed in real-time. An uninteresting DNA molecule located in a pore can be ejected by sending an “unblock” message back to the control software. This message leads the sequencer to reverse the voltage across the pore, causing the molecule to exit the pore in the reverse direction. The primary requirement for such a live depletion system is that the software making ejection decisions can keep up with the sequencing speed for up to 512 nanopores that concurrently sequence DNA molecules on a MinION sequencer. Two recent publications describe the implementation of such systems for specific settings. Payne et al. combined Oxford Nanopore’s Guppy base caller (8) with the minimap2 read aligner (11) in their Readfish workflow to make ejection decisions after mapping the reads to a reference genome in realtime. Kovaka et al. skipped the base-calling step and performed ejection decisions directly on nanopore current signals. While the latter is designed to run on a general-purpose CPU, it cannot handle large human size reference genomes. In contrast, Readfish can handle larger references but needs GPUs and an additional base-calling server for real-time base calling.

Furthermore, the usage of minimap2 (11) for read classification is not optimal. In their study, Payne et al. showed that only 83% of target reads were correctly classified for rejection after 0.8 seconds of sequencing. Marquet et al. observed the same issue when they tried to deplete all human host reads from vaginal samples with ONT’s adaptive sampling option. Using the depletion method supported by MinKNOW, 25% of human reads could not accurately be rejected by the software, leaving many reads that also pose ethical problems as patient consent is required for scientific data sharing. Further, missed mappings to repetitive regions of the reference genome can lead to delayed classifications when longer parts of the DNA molecule must be sequenced to make a rejection decision. Both lower sensitivity and classification delay will cause decreased enrichment of clinically relevant sequences of undetected pathogens or antibiotic resistance markers.

This study introduces *ReadBouncer* as a new tool for nanopore adaptive sampling that combines state-of-the-art base-calling software with the DREAM index (14, 15) Read-bouncer facilitates both GPU base-calling with ONTs Guppy as well as CPU base-calling with DeepNano-blitz (16). Its Interleaved Bloom Filter (IBF) data structure allows for fast querying of hashed k-mers on large sequence datasets resulting in an improved read classification strategy. Within an integrated workflow, Readbouncer uses IBFs to classify base-called DNA fragments for ejection and finally communicates the decision to the sequencing control software.

We first investigate our read classification approach by comparing it to other software tools used for read classification in a nanopore adaptive sampling context. ReadBouncer shows the best accuracy, recall, F1-Score, and Matthews correlation coefficient (MCC) among all tools on a simulated and a real-world dataset, while having almost the same precision and specificity as the best competitor. Furthermore, our tool also has the smallest reference sequence index size and peak memory usage.

We also compare ReadBouncer with Readfish and ONT’s MinKNOW software using a playback run of a whole human genome sequencing experiment to evaluate its adaptive sampling performance. In this comparison, we demonstrate that ReadBouncer outperforms the other tools in a targeted sequencing experiment. ReadBouncer results consistently show more sequenced bases for target references and significantly shorter mean read length of off-target or rejected nanopore reads. These results indicate that ReadBouncer can make faster and more reliable rejection decisions than Readfish and MinKNOW. ReadBouncer’s source code and installer files for Windows and Linux are freely available as a Git repository (https://gitlab.com/dacs-hpi/readbouncer) under GNU General Public License 3 (GPL-3.0).

## Methods

### Read Classification

With the current nanopore sequencing speed of 450 nucleotides per second, an adaptive sampling approach ideally makes ejection decisions within 2 seconds after sequencing of a DNA molecule has started. This requires fast base calling and rapid and reliable classification of read fragments smaller than 500 nucleotides. Readfish (10) uses the long-read alignment tool minimap2 (11) for this purpose. Although being fast and accurate for long error-prone nanopore reads, the alignment approach poses some challenges when working with short error-prone fragments of less than 500 nucleotides. For optimal enrichment of low abundance genomic regions, we need to make reliable rejection decisions as fast as possible. Payne et al. showed in their study that it takes about 360 nucleotides for minimap2 to align 90% of those reads correctly. That means, if we want to get higher enrichment, we need to improve the classification sensitivity for the same read length. Mappings are also hard to use when there is no good quality reference sequence available for an organism that is the depletion target such as non-model organisms. In such scenarios, one would try to use the reference sequence of a closely related species for read classification. Mapping reads to the reference of a closely related species would fail to find numerous reads that we would aim to eject from the pore.

All these findings motivated us to seek a different, fast classification strategy. To our knowledge, the fastest current sequence comparison algorithms use k-mer based approaches, where a DNA sequence is divided into small overlapping substrings of size *k*. One approach, known as MinHash (17, 18), computes a hash value for every k-mer of a sequence and stores the smallest hash values within a data structure called a sketch. The same procedure is applied to the second sequence, and the number of hash values present in both sketches gives an accurate approximation of the identity between the two sequences. Although this works well for sequences of similar size, it fails for sequence containment tests, where one sequence is much smaller than the other one, which is the case when we want to check if a nanopore read is part of a reference genome.

A better approach for testing if the set of k-mers of a reference genome contains the k-mers of a read is using Bloom Filters (19, 20). A Bloom Filter simply is a bitvector of size n and a set of h independent hash functions. To insert a k-mer into a Bloom Filter, the bit positions that correspond to the *h* hash values of the k-mer are set to 1, and a k-mer is considered present in the Bloom Filter if all *h* positions return a 1 during the lookup phase. In our case, we would insert all k-mers of a reference genome into the Bloom Filter and lookup for the k-mers of a nanopore read in that Bloom Filter.

The biggest problem of k-mer based approaches is choosing the correct parameter value for *k*, which is always a tradeoff between sensitivity and specificity in the presence of sequencing errors. Larger values for *k* will result in more specific read classification results but will also fail to find many reads from the reference genome when the number of sequencing errors is high. When trying to classify nanopore reads with error rates of about 10%, the value for *k* will hardly become bigger than 13. The number of different k-mers of size 13 is combinatorially defined by 4^13^ = 67,108,864, which is much too small when working with human-sized genomes that compose about 3 billion k-mers of size 13. To overcome this issue, we divide the reference genome into overlapping fragments of size *m* and construct a separate Bloom Filter for each fragment. However, querying one read against each of the Bloom Filters separately reduces the performance of the Bloom Filter approach. Thus, we decided to use Interleaved Bloom Filters (IBF) as proposed by (14) to index the reference genomes.

An Interleaved Bloom Filter combines several Bloom Filters (bins) in one single bitvector. The IBF can be divided into several subvectors, each having the size of the number of bins. Since one bin in the IBF corresponds to one fragment of the reference sequence, the size of each subvector corresponds to the number of fragments. In Figure, for example, we divided the reference sequence into three overlapping fragments, each corresponding to one bin of the IBF. Thus, each subvector in the IBF consists of 3 bits. The *i*-th bit of every subvector belongs to the Bloom Filter bin of fragment *F*_*i*_. When inserting a k-mer from fragment *F*_*i*_ into the IBF, we compute all *h* hash values, which point us to the corresponding subvectors *SV*_*j*_ and then simply set the *i*-th bit of this subvector to 1.

When querying a read *p* against the IBF in order to check if it maps to any of the fragments, every k-mer of that read is matched against the IBF. That means we first retrieve the *h* subvectors *SV*_*j*_ and apply a logical AND to them, resulting in the required binning bitvector indicating the membership of the k-mers in the bins. The example in Figure visualizes this process. Here, the read consists of four 7-mers, for which we have to calculate the three hash values that point us to the corresponding subvectors *SV*_*j*_, as can be seen in particular for the 7-mer *CAGGATT*. A logical AND of these three subvectors gives us the binning bitvector for that 7-mer. In our example, the binning vector 010 for *CAGGATT* tells us that this 7-mer only matches fragment *F*_2_. Applying this procedure to every 7-mer of the read gives us four binning bit vectors. Finally, we only need to sum up the 1-bits in the binning vectors for every fragment, which gives us the number of matching 7-mers of the read for every fragment. Thus, instead of computing *h* hash values for every Bloom Filter separately, we only need to compute the *h* hash values once, which poses a significant reduction in computing time to investigate the membership of a k-mer in every Bloom Filter. This method enables us to quickly count the number of matching k-mers between the reference genome and a specific nanopore read. The challenge is to define a threshold value for the number of matching k-mers required to accept a certain nanopore read as a match against a fragment and thus as a match with the reference genome. In our example in Figure, we consider the read matching fragment *F*_2_ because three of the four 7-mers match with that fragment. In general, the best threshold value depends on the length of the nanopore read and the expected sequencing error rate. We will describe our method for determining this value in the next section.

### Optimal Bitvector Size

In a first step, ReadBouncer produces overlapping fragments of the given reference sequences, e.g., 100,000 nucleotide long fragments with an overlap of 500 base pairs. Each of those fragments represents a single bin in the Interleaved Bloom Filter. The constituting k-mers of each fragment are hashed using three different hash functions, and the bits of the corresponding index positions in the Interleaved Bloom Filter are set to one (Figure 1). Then, ReadBouncer automatically calculates the optimal IBF size in bits (*Bits*_*IBF*_) based on the following equations.

**Fig. 1.**
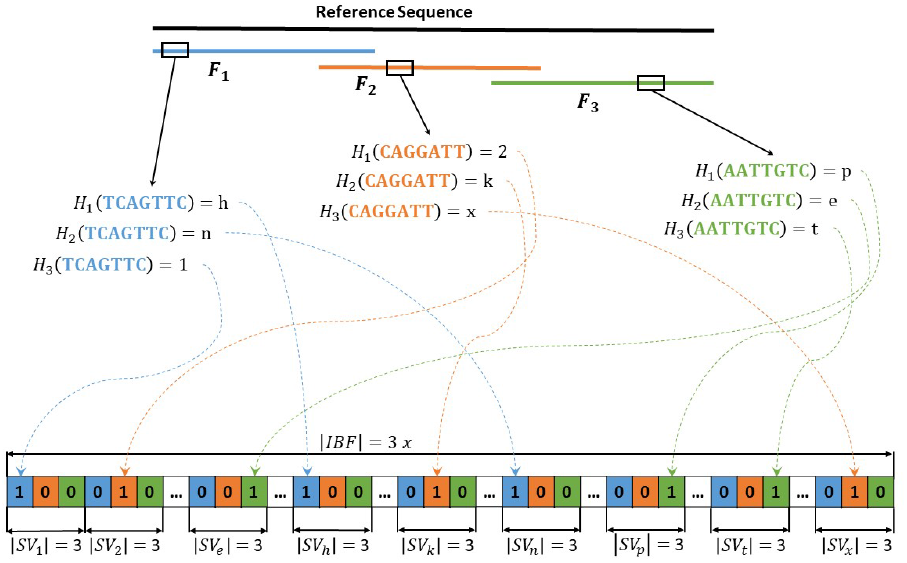
Example of an Interleaved Bloom Filter (IBF) construction. In the first step, we subdivide the reference sequence into three overlapping fragments. Then, for each k-mer of the differently colored fragments, all three hash values have to be calculated. The resulting hash values determine the subvector ^*SV*^*j* in which the corresponding bit is set to 1. For example, the second hash function for k-mer *AATTGTC* from fragment *F*3 returns *e*. Hence, we set the third bit of subvector *SVe* to 1. In this way, the three Bloom Filters for the three fragments are combined in an interleaved fashion. Since we have three fragments in our example, the length of every subvector is three, and the length of the IBF is 3*x*, where *x* is the defined length for every Bloom Filter of the three fragments.

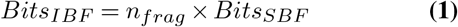

Where *n*_*frag*_ is defined as the number of fragments with maximum size *F* and *Bits*_*SBF*_ as a single Bloom filter size for a single fragment. Let *max*_*kmer*_ be the maximum number of k-mers for a fragment of size *F*, and k-mer size *k* be defined as

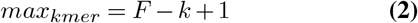

To calculate the optimal size for the IBF, we use the formula for finding the false positive rate in an IBF as proposed by (14).

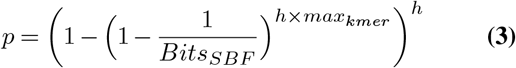

Then the optimal size of a single Bloom filter can be calculated by resolving the formula for *Bits*_*SBF*_ :

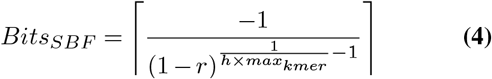

Where 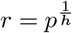, *h* is the number of used hash functions and *p* a predefined false positive rate. ReadBouncer implicitly uses three hash functions and a maximum false positive rate of 0.01 to minimize the number of false matches between the query sequence and a single bin of the Interleaved Bloom Filter.

### Minimum number of k-mer matches

During the read classification step, the k-mers of every read are hashed with the same three hash functions, and the number of matching k-mers for every bin is calculated as visualized in Figure 2. We accept a read as part of the reference sequence if the number of matching k-mers is greater than or equal to a given threshold *t* for at least one bin. We calculate the threshold using the expected sequencing error rate *e* and the definition of a (1 − *α*) confidence interval of the number of erroneous k-mers as recently provided by (21). They first defined the expected number of erroneous k-mers as follows:

**Fig. 2.**
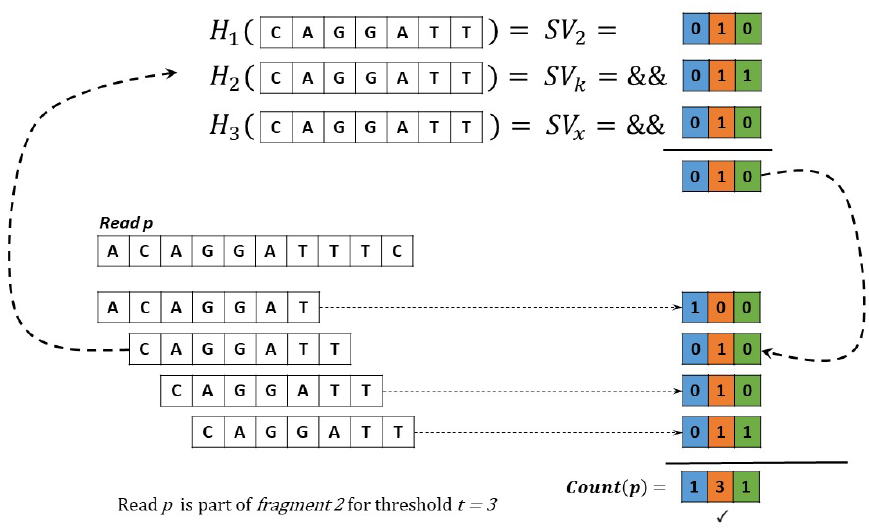
Finding the correct fragment for a given read p. For each k-mer of read p, we calculate the three hash values using the same hash functions as for the IBF construction. We use the resulting hash values to find the corresponding subvectors of the IBF. The sub bitvectors are combined with a bitwise AND to a binning bitvector. For all set bits in the binning vectors of the k-mers, we increment the counter of the corresponding bin in a counting vector. Bins whose counter is greater than or equal to a given threshold t are considered to contain the read p. In this example, we show the calculation of the binning bitvector for the 7-mer CAGGATT. Using the same three hash functions as for the IBF construction in Figure 2, we get the sub-vectors SV2, SVk, and SVx. We combine these three subvectors via logical AND to get the binning bitvector. The same procedure is applied to the other three 7-mers, and with the resulting four binning bitvectors, we can calculate the number of matching 7-mers of read p with each fragment. If at least three 7-mers match against one fragment, we accept the read as a match with the reference sequence.

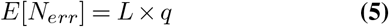

For a given read *r* with length *len*(*r*) and k-mer length *k*, we denote the number of k-mers of read *r* as *L* = *len*(*r*) − *k* + 1, and *q* is defined by (1 − (1 − *e*)^*k*^). In a second step, they show that the variance for the number of erroneous k-mers can be calculated by

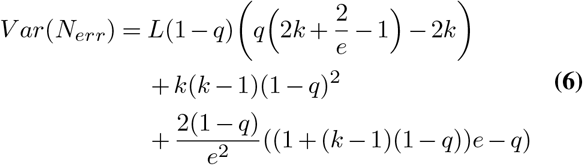

Finally, they define the (1 − *α*) confidence interval by:

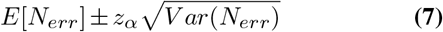

With 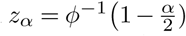, where we denote *ϕ*^−1^ as the inverse of the cumulative distribution function of the standard Gaussian distribution. Based on the calculation of the confidence interval for the number of erroneous k-mers, we define our threshold for the minimum number of matching k-mers for read *r* as:

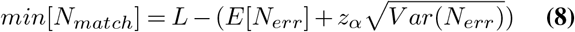

We classify a read as a match if the number of matching k-mers is bigger or equal to *min*[*N*_*match*_] for at least one bin in the IBF. ReadBouncer per default calculates this threshold for a 95%-confidence-interval, an expected sequencing error rate of 10%, and k-mer length 13. However, these values are adjustable via configuration parameters of the command line or graphical user interface (GUI).

### Workflow

The workflow of our tool consists of two consecutive parts. First, we build one or more indexes of the given reference sequence data set, which can be used as target or depletion filters. These indexes can be used directly in the second part of the workflow or stored on the computer hard disk for later usage. The construction of this index, for which we apply Interleaved Bloom Filters, is explained in further detail in section *Read Classification*.

The second part of our tool is the live-depletion or target-enrichment task (Suppl. Mat. Figure S2) Here, Readbouncer initially loads the indexes and waits for the nanopore device to start sequencing. Immediately after sequencing has begun, the sequencer streams raw electrical currents for every single molecule from every single sequencing pore of the flow cell to our integrated Read-Until client. Oxford Nanopore provides this functionality via an Application Programming Interface (API) of its MinKNOW control software (https://github.com/nanoporetech/minknow_api), which allows our Read-Until client to receive the raw data while the molecule traverses through the pore. The client is implemented in C++ and communicates with the MinKNOW control software via gRPC remote procedure calls (https://github.com/grpc/grpc).

Received raw signal data get pushed onto a base-calling queue, and a separate thread takes each reads raw signals from the queue and sends it to the chosen base-calling algorithm, which translates the electrical currents into a nucleotide string. The user can choose GPU base-calling with ONT’s Guppy basecaller for which we integrated a guppy client that communicates with a guppy basecall-server. Additionally, we integrated DeepNano-blitz (16) for the base-calling step, which is fast enough to perform the base-calling in real-time, even on CPUs.

Base called reads get pushed to the classification queue if the read length is bigger than or equal to 200 nucleotides, and another thread takes each read from that queue and passes it to the classification framework. Otherwise, the thread marks this read as “pending” and waits for the following data chunk to be base called and concatenates the base called sequences of the read until the minimum read length has been reached. The minimum read length of 200 nucleotides ensures higher confidence in the classification of the reads. In practice, this read length requirement will lead to most reads having about 360 nucleotides length, which corresponds to two data chunks sent by the MinKNOW software. The read classification thread then queries the read sequence against the loaded Interleaved Bloom Filter indexes as described in more detail in section *Read Classification*. Based on the classification, reads can either be marked for a rejection or continue further sequencing. If a read was not classified for rejection on a first try, we mark it as *once_seen* and wait for further sequencing data to try further classification attempts of that read. After the read has reached a maximum read length of 1,500 bp we stop trying to make ejection decisions and mark the read for continued sequencing as usual. Reads that have been classified for rejection or continued sequencing are finally pushed to the response queue and no further data chunks of that read are sent by the control software.

The last thread takes the classified reads from the response queue, and our Read-Until client sends response messages back to the MinKNOW control software for each read. The client sends an unblock message for reads that could be matched to the Interleaved Bloom filter, telling the sequencer to eject the corresponding DNA molecule. A *stop_further_data* message is sent to the control software for reads that were not classified for rejection. This message tells MinKNOW to continue sequencing the corresponding DNA molecule and send no additional chunks of data for that read.

## Results

In this study, we show how adaptive sampling benefits from our improved read classification approach. Therefore, we designed experiments that specifically focus on the evaluation of this approach when applied to both adaptive sampling strategies, depletion and targeted sequencing. In a first step, we compare ReadBouncer to minimap2 (11), which is used for classification by Readfish, and the pan-genomics matching tool SPUMONI (22), which is proposed as an alternative to minimap2 in targeted nanopore sequencing pipelines. Here, we assess all three tools on simulated and real reads from a recently published microbial mock community (23). In a second experiment, we compare ReadBouncer with Readfish in an adaptive sampling setting using the playback feature offered by Oxford Nanopore’s MinKNOW software to replay an already completed sequencing run. We assess both tools by targeting chromosomes 21 and 22 in a human whole genome sequencing run, looking at their ability to correctly filter out all other human nanopore reads. Here, we do not compare against SPUMONI because there exists no adaptive sampling pipeline integrating SPUMONI for read classification.

We perform all experiments for classification performance assessment on a laptop with a 2.8 GHz Intel Core i7-7700HQ CPU and 16 GB of memory with an Ubuntu 20.04 OS installed. For the classification evaluation, We run each tool with a single thread for runtime comparisons and record the wall clock time and peak resident set size (RSS) reported by the individual tools or GNU time 1.7.

### Evaluating Read Classification

#### Experimental Setup

During a nanopore targeted sequencing experiment with ONT’s ReadUntil functionality, the sequencing device transmits electrical current data via the MinKNOW control software to ReadBouncer. This data is received as chunks, representing a maximum of 0.4 or 0.8 seconds of sequencing, depending on the MinKNOW configuration. Since a DNA molecule translocates through the pore at a speed of about 450 bases per second, 0.4 seconds of sequencing represent about 180 bases of data. In the following experiments, we mimic the situation where a chunk represents 0.4 seconds of sequencing data received and base called immediately by an adaptive sampling tool. Since we aim to make rejection decisions as early as possible while still being able to classify most of the reads correctly, we want to assess the classification accuracy of the three tools after two chunks of data, which correspond to 360 nucleotides or 0.8 seconds of sequencing. In this section, we assume that base-calling has already been performed. For a fair comparison, we set up all experiments in such a way that all three tools, minimap2, SPUMONI, and ReadBouncer, attempt to classify reads based on the 360 bases long read prefix. In practice, all reads, both simulated and real reads, were cut to only the first 360 bases. ReadBouncer then hashes all k-mers of these 360 bases and compares the hash values to a prebuilt Interleaved Bloom Filter of the depletion target references to make classification decisions.

We use the software’s default settings for the SPUMONI approach, which means splitting the prefix into substrings of 90 nucleotides each for further read classification. SPUMONI also needs a prebuild index of the references but has to include the reverse complement of the depletion target references. SPUMONI matches the substrings against this positive index and a null index, consisting of the reverse sequences of the positive index. Finally, classification decisions are made by using a Kolmogorov-Smirnov test.

For the minimap2-based approach, we mimic the read classification of Readfish by using the mappy Python interface (https://pypi.org/project/mappy/) for minimap2. For comparison, we align the read prefixes with the *map-ont* settings without applying any mapping quality filter, which are the same settings used by Readfish. To evaluate the three tools reads correctly classified as belonging to the depletion target are considered true positives (TP), while reads falsely classified for depletion are called false positives (FP). Consistently, reads that are correctly not classified as depletion target are considered true negatives (TN), and reads belonging to the depletion target but not classified for depletion are called false negatives (FN). We calculate the classification accuracy, precision, recall, specificity, and F1-score for all three approaches based on those considerations. Since we assume an imbalanced number of sequenced reads between depletion and enrichment targets, we also report the Matthews correlation coefficient (MCC) in every experiment.

#### Simulated Mock Community

In the first dataset, we consider simulated ONT-like reads derived from the identical genomes of the ZymoBIOMICS High Molecular Weight DNA Mock Microbial community (ZymoMC) (23). This mock community consists of seven bacterial species - *Enterococcus faecalis, Listeria monocytogenes, Bacillus subtilis, Salmonella enterica, Escherichia coli, Staphylococcus aureus*, and *Pseudomonas aeruginosa* – as well as *Saccharomyces cerevisiae*. We use PBSIM2 (24) to simulate Oxford-Nanopore-like reads (R9.4 pores) from Zymo Mock Community references at varying levels of mean read accuracy: 80%, 85%, 90%, 95%, and 98%. Furthermore, we simulated proportions of reads from each genome in such a way to mimic a scenario where only 2.16% of reads originate from *Saccharomyces cerevisiae* (Suppl. Mat. Figure S3). The goal here is to enrich *Saccharomyces cerevisiae* sequences by correctly classifying bacterial reads, which we would aim to eject from the pores in a real nanopore sequencing run. This can be considered as a depletion-only experiment, where a priori only the depletion references are known, but not the enrichment targets. Therefore, we build an index of the seven bacterial reference genomes and query all bacterial and yeast reads against the index. Consistent with our definition in section *Experimental Setup* we consider correctly classified bacterial reads true positives (TP), while yeast reads found in the index are considered false positives (FP). In addition, we define bacterial reads that are missed to be found by a tool in the index as false negatives (FN), and yeast reads that are not found in the index are considered true negatives (TN).

On all read accuracy levels, ReadBouncer consistently demonstrates best accuracy, recall, precision, F1-scores, and MCC (Suppl. Mat. Table S5). In Figure 3, we visualize recall and specificity for the three tools across various read accuracies. It can be observed that recall improves with increasing read accuracy for all three tools while specificity stays almost unchanged. On all read accuracy levels, ReadBouncer demonstrates slightly but consistently better recall (sensitivity) than SPUMONI, while both tools outperform minimap2. Minimap2 is the only tool that shows 100% specificity, but ReadBouncer comes close to 100% as well. SPUMONI lags a bit behind the specificity scores of the other two tools. It can be seen that ReadBouncer is the best performing tool for this read classification task. It combines high recall (sensitivity) with high specificity. The other two tools either have high recall but lower specificity or high specificity but lower recall scores.

**Fig. 3.**
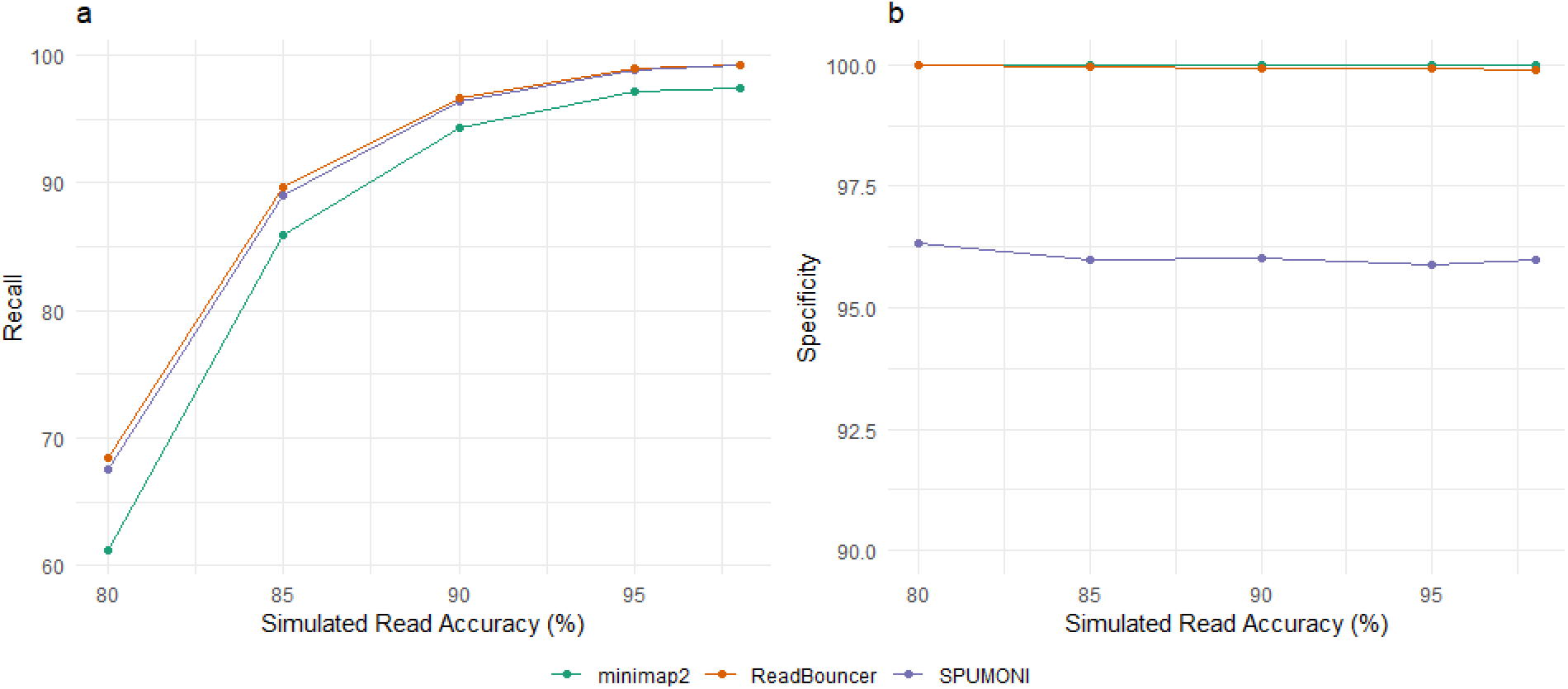
Visualization of *(a)* Recall and *(b)* Specificity with varying simulated read accuracies for ReadBouncer, minimap2, and SPUMONI.

#### Real Mock Community

Next, we applied our method to real nanopore reads from a Zymo Mock Community (NCBI BioProject PRJNA742838). After sample preparation, we sequenced the mock sample on a MinION flowcell (FLO-MIN106) with v.R9.4.1 pores (Suppl. Mat. Section S1). Obtained Fast5 files were base called with DeepNano-blitz using a recurrent neural network size of 48. For better comparison with minimap2, we first build a separately obtained minimap2 mapping as a gold standard. Therefore, we filter out all reads shorter than 2,000 base pairs and trim the first 360 nucleotides from each read since we use these bases for later classification. Then, we mapped the trimmed reads with standard ONT settings to the ZymoMC reference genomes and only reads with a mapping quality score bigger or equal to 30 are considered confidently mapped. From these mapped reads, the trimmed 360 nucleotide long prefixes are used for the read classification by the three tools again. Proportions of reads from each genome are similar to the simulated experiment with 2.27% of reads from *Saccharomyces cerevisiae* (Suppl. Mat. Figure S4). In this experiment, we also measure the peak Resident Set Size (RSS) and index size in GigaByte and the throughput for each of the tools in reads classified per second.

Results in Table 1 show that ReadBouncer achieves better accuracy, recall, and F1-score than SPUMONI and minimap2, which both have similar results for those three measures. Minimap2 has slightly better precision and specificity than ReadBouncer. While SPUMONI has almost the same precision as ReadBouncer and minimap2, it shows significantly less specificity. These results are consistent with those for the simulated data sets in section *Simulated Mock Community* and show that ReadBouncer outperforms the other tools on read classification for short nanopore reads.

**Table 1.** Comparing ReadBouncer, SPUMONI, and minimap2 across various metrics on a real Zymo Mock Community data set consisting of seven bacterial species and *Saccharomyces cerevisiae*. Reads from a nanopore sequencing run are mapped to the eight organisms to generate ground truth. We use only the first 360 nucleotides for classification from those confidently mapped reads to mimic unblock decision-making after 0.8 seconds of sequencing the individual read. All reads are mapped against the seven bacterial reference sequences to filter out only the bacterial reads. At the same time, we want to keep as much *Saccharomyces cerevisiae* reads, which corresponds to an enrichment of that organism in an enrichment/depletion experiment. Consistent with the simulated data, ReadBouncer can classify a higher percentage of bacterial reads while having only slightly less precision and specificity than minimap2. Our approach also shows to be the computationally most effective one, with the lowest memory footprint and highest classification throughput.

| Tool | ReadBouncer | SPUMONI | minimap2 |
| --- | --- | --- | --- |
| Accuracy | **94.50** | 90.96 | 89.33 |
| Precision | 99.99 | 99.89 | **100.00** |
| Recall | **94.38** | 90.85 | 89.08 |
| Specificity | 99.73 | 95.87 | **99.95** |
| F1-Score | **97.10** | 95.16 | 94.23 |
| MCC | **0.52** | 0.41 | 0.39 |
| Peak RSS (GB) | **0.099** | 0.163 | 0.272 |
| Index Size (GB) | **0.047** | 0.153 | 0.097 |
| Throughput (reads per sec) | **5967** | 1102 | 5632 |

Another important aspect is the amount of main memory a tool needs to hold the reference index needed for read classification. Using the seven bacterial reference genomes of the Zymo Mock Community as depletion target (reference index), ReadBouncer shows the smallest maximum memory consumption measured as Peak Resident Set Size (RSS). It only needs 0.099 GigaBytes (GB) of main memory, in contrast to 0.272 GB consumed by minimap2. Furthermore, ReadBouncer has the smallest index file size (0.047 GB) of all three tools. In addition to the smallest memory footprint, ReadBouncer also achieves the highest classification throughput. We can classify 5967 reads per second with our approach compared to 5632 reads per second by minimap2 and 1102 reads per second achieved by SPUMONI. These results show that we cannot only correctly classify more reads than the other tools. ReadBouncer is also computationally more efficient than other state-of-the-art read classification tools used for nanopore adaptive sampling.

#### Adaptive Sampling Evaluation

In our live experiment, we assess our read classification based on Interleaved Bloom Filters in a targeted adaptive sampling setup. For this purpose, we downloaded a bulk FAST5 file of a human whole-genome sequencing experiment provided via the Github page of Readfish (https://github.com/LooseLab/readfish). Such a bulk FAST5 file (25) allows the playback of the whole sequencing run for testing if the ReadUntil functionality is working correctly. Oxford Nanopore’s MinKNOW software simulates an already finished sequencing run without the need for a physical sequencing device when performing a playback run. Compared to the original sequencing run, read signals are reported at the same time point after starting the run. Unblocking a read does not cause MinKNOW to finish sending signals for that read during a playback run. It just breaks the read when receiving an unblock message for the read and creates a new read identifier but continues to send signals of the same original read. Here, we compare ReadBouncer and Readfish using both real-time GPU base-calling with ONT’s Guppy basecaller and real-time CPU base-calling with DeepNano-blitz. Both tools were run on our Ubuntu Laptop while GPU live base calling was performed on an NVIDIA Jetson AGX Xavier. Additionally, we compared the results with two MinKNOW adaptive sampling experiments, one using MinKNOW’s *target* and the other using MinKNOW’s *deplete* method. Both experiments were performed on the NVIDIA Jetson AGX Xavier, too.

In our experiments, we do a playback of a complete human genome sequencing run with the goal to enrich for chromosomes 21 and 22 of the human genome and deplete all other human reads from that run. This setup not only mimics a targeted sequencing approach, it is also corresponds to the application of sequencing a clinical human blood sample where up to 99% of the reads are human reads that we would want to deplete in order to enrich the number of reads from a pathogenic microbe. We perform playback runs for 60 minutes on ONT’s MinKNOW control software version 4.3.3. To ensure that the vast majority of the sequenced reads are of human origin, we first perform a playback run without adaptive sampling. Reads were base called with Guppy 5.0.14 and mapped with minimap2 to the human Telomere-to-Telomere Consortium (“T2T”) CHM13 v1.1 reference assembly (26). From the resulting reads passing the in-built quality filtering of MinKNOW, 99.66% could be mapped to the human reference genome. For the comparison of the tools in an adaptive sampling seeting, we first adjust the *break_reads_after_seconds* parameter within MinKNOW to 0.4 seconds as recommended by the Readfish authors. Since MinKNOW sends data as chunks, this parameter sets the size of one chunk to a maximum of 180 nucleotides. Both tools, ReadBouncer and Readfish, can concatenate the data chunks and perform classification after the receipt of every chunk. For integrated CPU base calling with DeepNano-blitz we used a neural network size of 48 for both tools. For real-time GPU base calling on the NVIDIA Jetson AGX Xavier we used the fast base-calling mode of Guppy 5.0.14 basecall server for ReadBouncer and Readfish. For the evaluation of both tools, we repeat the same play-back run for 60 minutes. In the experiment with CPU basecalling, we ran ReadBouncer with default parameters (*fragment*_*size* = 100, 000 and *kmer*_*size* = 13) using three base calling threads and three read classification threads, respectively. The same setting was applied to Readfish with three CPU basecalling threads and minimap2 using three threads per default. Since Guppy ensures a higher raw read accuracy, we ran ReadBouncer with *fragment*_*size* = 200, 000, *kmer*_*size* = 15 and *error*_*r*_*ate* = 0.05 in the GPU basecalling experiment. In all experiments, we used chromosomes 21 and 22 as target filter and all other chromosomes as depletion filter in ReadBouncer. Our settings within the Readfish configuration file correspond to the example TOML file in the github repository (https://github.com/LooseLab/readfish/blob/master/examples/human_chr_selection.toml) and aim to target chromosomes 21 and 22 as well, while unblocking all reads that do not map to the targets. For MinKNOW *target* we used chromosomes 21 and 22 as reference and for MinKNOW *deplete* all other chromosomes as reference sequence.

After finishing the playback run, the resulting fast5 files were basecalled in high accuracy mode with Guppy 5.0.14. All reads in the resulting fastq files were mapped to the human genome reference and mapping statistics were calculated with Readfish’s *summary* script.

The results of all six experiments can be seen in Table 2. Our first observation is that the results for our target chromosomes 21 and 22 are similar for all experiments but the MinKNOW *deplete* experiment. Here, the number of on-target reads is much higher while showing the smallest mean and median read lengths caused by a high number of false rejection decisions. These results suggest that MinKNOW *deplete* is not suitable for targeting single chromosomes of the human genome in an adaptive sampling experiment. On the other hand, MinKNOW *target* shows similar results for chromosomes 21 and 22 when compared to ReadBouncer and Readfish. However, the mean read length of 3,629 bp measured for unblocked reads is much higher than those in the ReadBouncer and Readfish experiments, which shows that MinKNOW *target* spends too much time sequencing off-target reads. These experiments show that ReadBouncer and Readfish outperform the two MinKNOW adaptive sampling strategies.

**Table 2.**
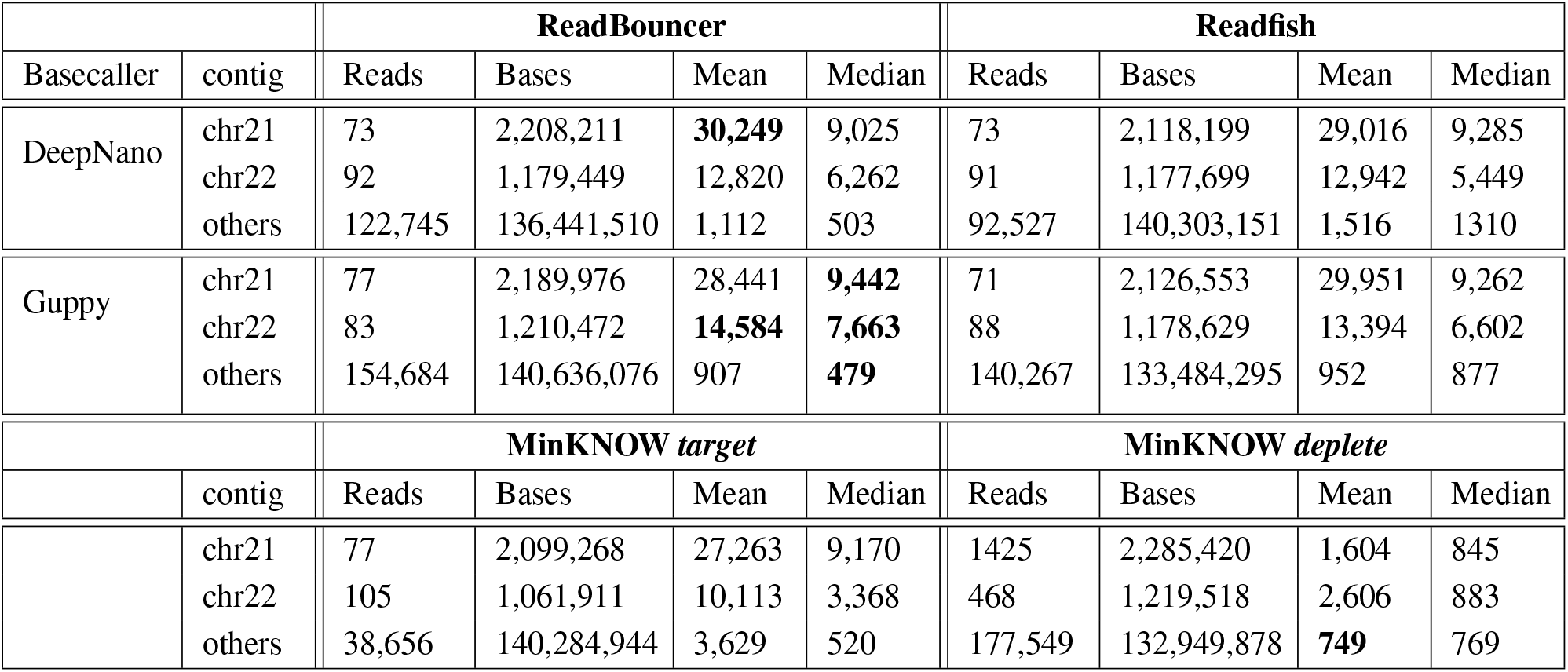
Comparison of ReadBouncer, Readfish and MinKNOW in a targeted sequencing experiment. Four 60 minute playback runs of a whole human genome sequencing experiment were performed using either ReadBouncer or Readfish in combination with either DeepNano CPU base calling or Guppy GPU base calling. The same experiment was repeated with MinKNOW’s adaptive sampling functionality in *target* and *deplete* mode. The goal of all experiments was to target chromosomes 21 and 22 while rejecting all other human reads. ReadBouncer and Readfish show consistently better results when using GPU base calling with ReadBouncer having shorter mean and median read lengths for non-target reads regardless of the used basecaller. ReadBouncer outperforms MinKNOW *target* by having longer read lengths for on-target reads and shorter read lengths for off-target reads caused by a better read classification. MinKNOW *deplete* has the worst results of all tools indicated by high numbers of on-target reads with short read lengths caused lots of false unblock decisions for on-target reads.

|  |  | ReadBouncer |  |  |  | Readfish |  |  |  |
| --- | --- | --- | --- | --- | --- | --- | --- | --- | --- |
| Basecaller | contig | Reads | Bases | Mean | Median | Reads | Bases | Mean | Median |
| DeepNano | chr21 | 73 | 2,208,211 | **30,249** | 9,025 | 73 | 2,118,199 | 29,016 | 9,285 |
|  | chr22 | 92 | 1,179,449 | 12,820 | 6,262 | 91 | 1,177,699 | 12,942 | 5,449 |
|  | others | 122,745 | 136,441,510 | 1,112 | 503 | 92,527 | 140,303,151 | 1,516 | 1310 |
| Guppy | chr21 | 77 | 2,189,976 | 28,441 | **9,442** | 71 | 2,126,553 | 29,951 | 9,262 |
|  | chr22 | 83 | 1,210,472 | **14,584** | **7,663** | 88 | 1,178,629 | 13,394 | 6,602 |
|  | others | 154,684 | 140,636,076 | 907 | **479** | 140,267 | 133,484,295 | 952 | 877 |
|  |  | MinKNOW *target* |  |  |  | MinKNOW *deplete* |  |  |  |
|  | contig | Reads | Bases | Mean | Median | Reads | Bases | Mean | Median |
|  | chr21 | 77 | 2,099,268 | 27,263 | 9,170 | 1425 | 2,285,420 | 1,604 | 845 |
|  | chr22 | 105 | 1,061,911 | 10,113 | 3,368 | 468 | 1,219,518 | 2,606 | 883 |
|  | others | 38,656 | 140,284,944 | 3,629 | 520 | 177,549 | 132,949,878 | **749** | 769 |

Comparing ReadBouncer with Readfish, when both tools use the same basecaller, ReadBouncer shows slightly better results regarding median read lengths and the number of bases sequenced. We also see that the choice of the base calling tool has a significant impact on the outcome of the adaptive sampling experiment. Using Guppy GPU base calling for both tools, ReadBouncer and Readfish results in much shorter read read lengths for non-target (unblocked) reads. Interestingly, we observe that unblocked reads from the ReadBouncer playback runs have shorter mean and median read lengths than those from the Readfish playback runs. This is also shown in the length distribution plots of unblocked reads for playback runs with Guppy base calling presented in Figure 4.

**Fig. 4.**
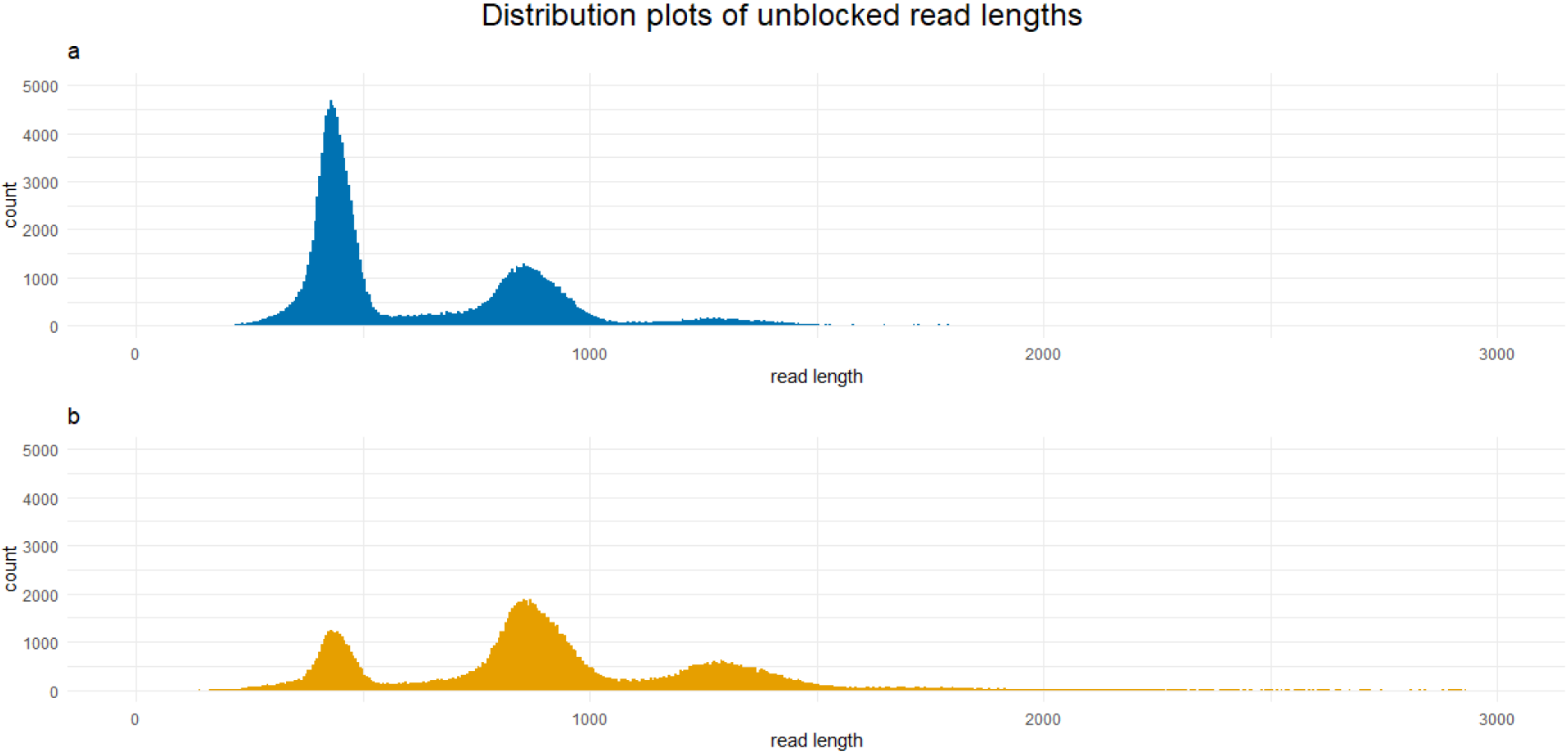
Read length distributions of unblocked reads when using *a)* ReadBouncer or *b)* Readfish on a 60 minutes playback run of a whole human sequencing experiment with real-time Guppy GPU base calling. ReadBouncer makes faster rejection decisions than Readfish, which can be observed by shorter read lengths of unblocked nanopore reads.

## Discussion

The idea of adaptive sampling is to selectively sequence individual DNA molecules on nanopore sequencing devices using in-silico methods. This study presents a new tool for adaptive sampling that improves read classification by combining interleaved Bloom filters with k-mer matching statistics. ReadBouncer shows a higher read classification sensitivity than other state-of-the-art classification tools for adaptive sampling while retaining a high specificity. Our tool also improves classification performance and memory usage compared to the other tools. We could observe much shorter read lengths of non-target reads in different playback experiments when using ReadBouncer instead of Readfish. This means that we can investigate more DNA molecules in the same amount of sequencing time with ReadBouncer. We developed our tool as an easy-to-install software application with a graphical user interface on Linux and Windows operating systems. Additionally, ReadBouncer supports fast CPU base-calling, providing even small sequencing facilities or in-field researchers that typically only have access to low-cost hardware the possibility to use the adaptive sampling feature of the MinION sequencer.

The key benefit of our new tool is the improved read classification. We neither use signal nor sequence space mapping algorithms for read classification compared to other adaptive sampling tools. Instead, our Interleaved Bloom Filter approach uses k-mer counting in Bloom Filters for sequence containment testing, resulting in smaller index files and fewer memory requirements. However, the improved sensitivity comes at the cost of decreased classification speed with increasing reference database size due to our approach of fragmenting the reference genome sequences and using one bin of the Interleaved Bloom Filter per fragment. The fragmentation approach ensures a high classification specificity for nanopore reads with high error rates of approximately 10-15% as observed by the CPU basecaller DeepNano-blitz (16). This error rate forces us to use small k-mer sizes such as 13, which entails the need for smaller fragment sizes down to 100,000 nucleotides to avoid too many false positive matches. Using real-time GPU base-calling with single raw read accuracies of about 94% allows increasing the k-mer size to 15 and fragment size to 200,000, reducing the number of bins in the interleaved Bloom filter by 50%. In the future, we expect to use even fewer fragments per genome and consequentially improve the classification speed for larger genomes as Oxford Nanopore is steadily improving its per-read accuracy. This would also enable the usage of our Interleaved Bloom Filter approach for real-time metagenomics classification of nanopore reads or the construction of pan-genomics indexes that store all different haplotypes of a pathogen in one IBF, with one haplotype per bin. To further increase performance, combining ReadBouncer and minimap2 could be worthwhile, as the integration of different methods in related fields has demonstrated (27).

A second key feature of ReadBouncer is its support for fast and accurate real-time GPU base-calling with ONT’s Guppy and real-time CPU base-calling with DeepNano-blitz. This study showed that both approaches show reliable results for a whole human sequencing playback run with the application to enrich specific chromosomes while rejecting reads belonging to all other chromosomes. Since there are some performance drawbacks of MinKNOW when using a playback run, the measured read lengths of rejected reads can deviate to a real experiment. Other users (https://github.com/sirselim/jetson_nanopore_sequencing) reported much shorter unblocked read lengths on real experiments performed on NVIDIA Jetson AGX Xavier. To ensure reproducibility and fair comparison between tools and to reduce the influence of potential artifacts, we evaluated our tool here on a playback of a well-performed experimental run rather than during run-time of the sequencer. Since a playback run can be regarded as real data from a real sequencing experiment, we do not expect any bias resulting from this comparison. The improved sensitivity of ReadBouncer’s classification approach is also not influenced by the usage of a playback run.

To our knowledge, ReadBouncer is the first tool that officially supports CPU-based adaptive sampling on commodity hardware. In this study, we compared the combination of ReadBouncer and DeepNano-blitz with Readfish using DeepNano-blitz, showing far better results by ReadBouncer. Since CPU base-calling using DeepNano-blitz is not officially supported by Readfish, we assume that there are still some performance issues within Readfish that cause the big difference in unblocked read length between the two tools. Further improvements in Readfish could reduce this gap. However, we still expect Readbouncer to outperform Readfish on CPUs because of its better read classification.

We expect that ReadBouncer can make a significant contribution to the field of pathogen detection in non-model organisms. Metagenomics sequencing of such samples easily consists of up to 99% host reads that can be depleted with adaptive sampling. Here, our CPU-based approach also makes access to adaptive sampling much easier for researchers studying wild living animals in the field. With Nanopores being successfully applied to peptide sequencing (28), a possible extension of the approach may also be to targeted protein sequencing.

Another potential use case for adaptive sampling is the real-time detection of antibiotic resistance and virulence genes. In their recently published study, Zhou et al. showed that direct nanopore metagenomics sequencing of human blood samples could detect pathogens in real-time but fails to detect antibiotic resistance genes. They compared direct metagenomic sequencing approach to MinION sequencing of blood culture samples. Using blood cultures, they could deplete human reads to about 65% of all sequenced reads in the corresponding sample, which was sufficient to identify more than 80% of resistance genes after 2 hours of sequencing. We expect that the number of sequenced human host reads can be depleted at a much higher rate by using adaptive sampling. This could supersede the need for blood cultures in such clinical settings and reduce costs and decrease the time to detect pathogens in human blood samples. Furthermore, a point-of-care test for antibiotic resistance genes in human patient samples that also avoids shipping the samples to a nearby laboratory could massively decrease antibiotic drugs’ usage and help restrict the development of antibiotic resistance that are a burden to many health care systems all over the world. Besides further sample preparation and sequencing technology improvements, we encourage scientists to set up proof-of-principle studies investigating the potential application of adaptive sampling for real-time antimicrobial resistance gene detection.

## Acknowledgements

The authors thank Vitor C. Piro and Tobias P. Loka (HPI) and Knut Reinert (FU Berlin) for valuable discussions and comments on the usage of IBFs and Martin Beer (FLI) for insights into Nanopore sequencing. We thank the Genome Sequencing Unit at Robert Koch Institute for sequencing of the ZymoBIOMICS Mock Community.

