## Supplemental Material for "ReadBouncer: Precise and Scalable Adaptive Sampling for Nanopore Sequencing"

[*] To whom correspondence should be addressed

### S1 DNA Sequencing

ZymoBIOMICS HMW DNA Standard was purchased from Zymo Research, from which 400 ng were transferred into a 1.5 mL DNA LoBind tube, and the volume was adjusted to 7.5 µL with Nuclease-free water. The sample was mixed by flicking the tube, spun down briefly, and then transferred into a 0.2 mL PCR tube. 2.5 µL Fragmentation Mix was added, and the tube was mixed by flicking the tube and spun down. After incubating for 1 min at 30 °C and subsequently for 1 min at 80 °C, the tube was briefly put on ice. 1 µL RAP was added to the barcoded DNA.
After mixing by flicking the tube and spinning down, the sample was incubated for 5 min at room temperature. In a new DNA LoBind tube, 34 µL Sequencing Buffer (SQB), 25.5 µL Loading Beads (LB), 4.5 µL Nuclease-free water, and 11 µL DNA library were added. Meanwhile, for the priming of the flow cell FLO-MIN106 (R9.4 SpotON), 30 µL Flush Tether (FLT) was added directly to a tube of Flush Buffer (FB) and mixed by vortexing. 800 µL priming mix were loaded into the flow cell via priming port without introducing any air bubbles. After incubating for 5 min, the SpotON sample port was opened, and 200 µL priming mix were loaded into the flow cell via priming port. Finally, the prepared DNA library was mixed by pipetting up and down, and 75 µL of the sample volume was loaded into the flow cell via the SpotON sample port in a dropwise fashion. After closing the SpotON sample port and priming port and replacing the MinION lid, the sequencing experiment was started via MinKNOW software.

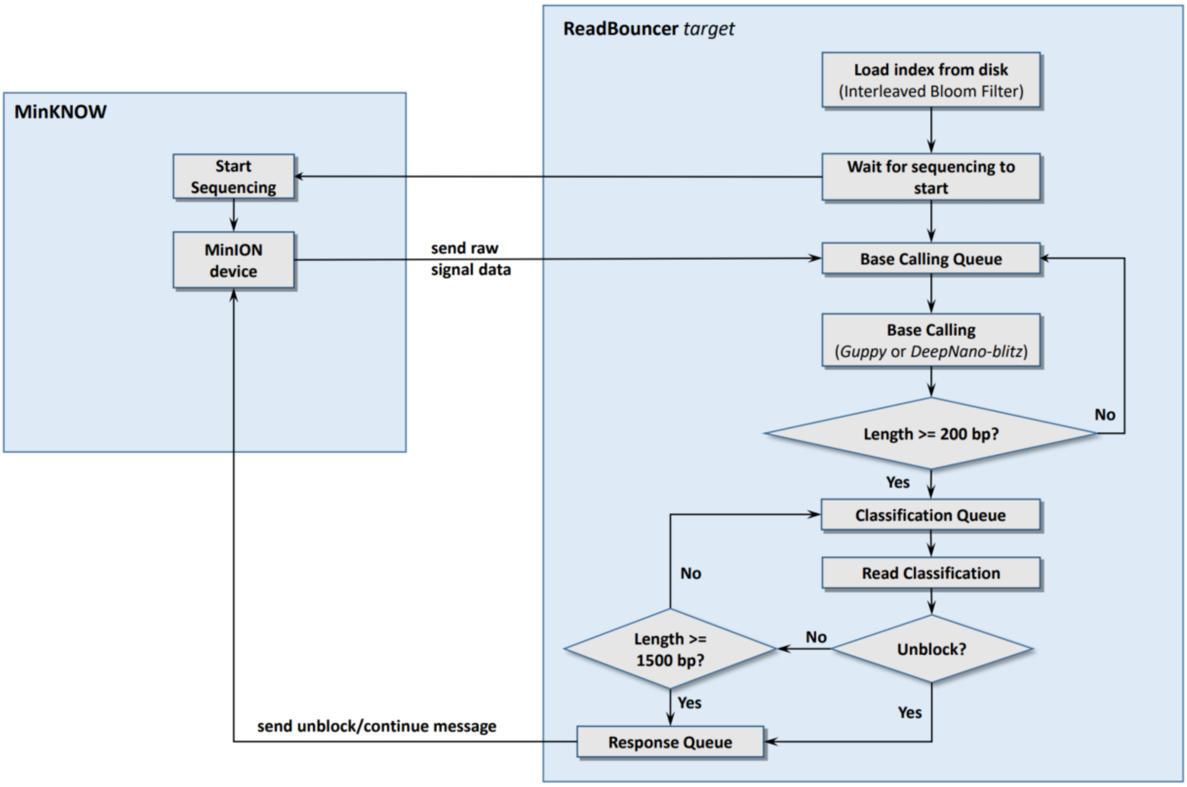


**Figure S2:** Flow diagram of ReadBouncer's adaptive sampling workflow. Interleaved Bloom Filter(s) of reference sequences are loaded from index file(s) first. ReadBouncer then waits until the MinKNOW starts the sequencing run. When sequencing has begun, the MinION device sends raw signals for every DNA molecule currently traversing a nanopore to ReadBouncer, which pushes the signals onto a base-calling queue. The base-calling thread takes signals from the queue and performs base calling via Guppy or DeepNano-blitz. The base-called sequences get pushed onto the classification queue if the length is equal to or longer than 200 nucleotides or pushed back to base calling queue otherwise. The classification thread takes sequences from the classification queue and queries the sequences against the depletion and/or target IBFs. If a sequence is found in the depletion IBF but not in the target IBF, the corresponding read is marked for unblocking and pushed onto the response queue. If the sequence is not found in the depletion IBF or is found in the target IBF and the sequence length is shorter than 1,500 bp, the corresponding read is pushed back to the classification queue. ReadBouncer repeats the classification procedure using consecutive chunks of data until the sequence length exceeds 1,500 bp. Reads that were not classified to reject are marked for sequencing as usual. A stop_further_data message for those reads is pushed onto the response queue. Finally, the response thread sends back action messages of reads from the response queue to the MinKNOW software and MinION device, respectively.

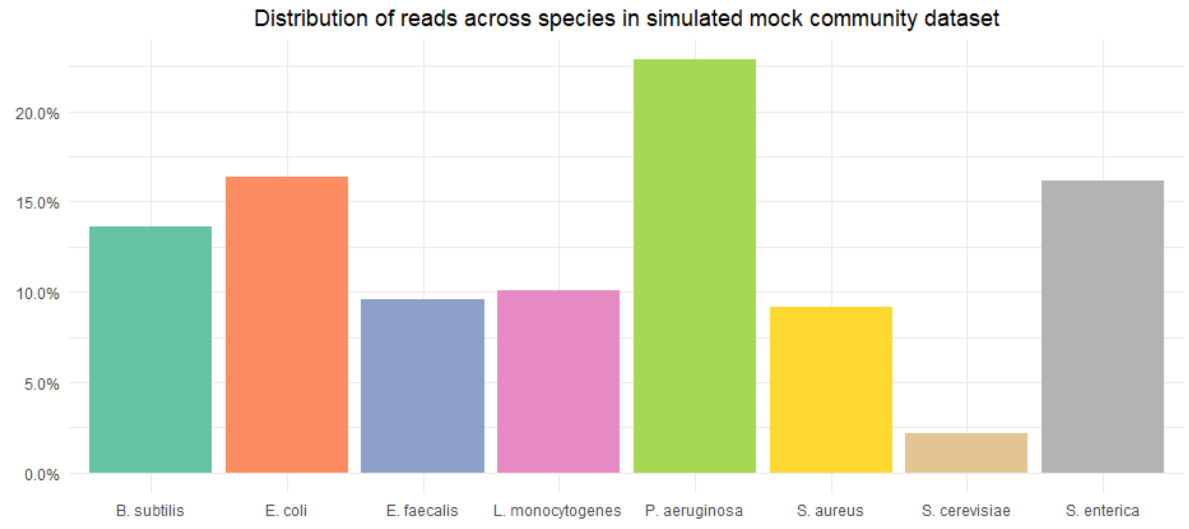


**Figure S3:** Proportion of reads from each species in the simulated mock community dataset. Less than 2.5% of reads were simulated from *Saccharomyces cerevisiae*. We would aim to deplete bacterial reads in order to enrich for *Saccharomyces cerevisiae*.

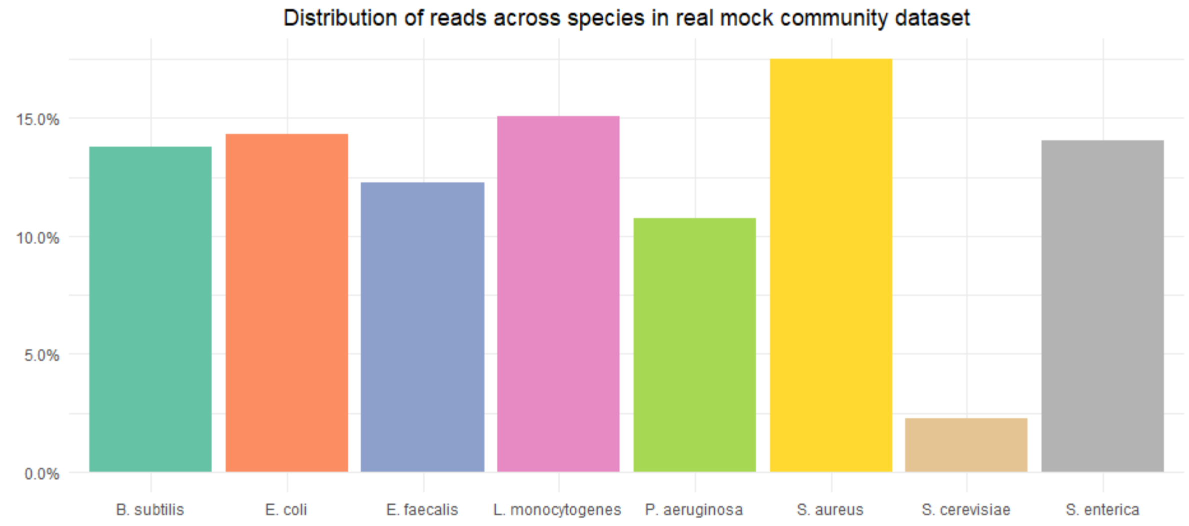


**Figure S4:** Proportion of reads from each species in the real mock community dataset. Only 2.5% of reads sequenced from a real mock community originate from *Saccharomyces cerevisiae*. We would aim to deplete bacterial reads in order to enrich for *Saccharomyces cerevisiae*.

| Read Accuracy(%) | 80 | | | 85 | | | 90 | | |
|---|---|---|---|---|---|---|---|---|---|
| **Tool** | **Read-Bouncer** | **SPUMONI** | **minimap2** | **Read-Bouncer** | **SPUMONI** | **minimap2** | **Read-Bouncer** | **SPUMONI** | **minimap2** |
| Accuracy | **69.16** | 68.23 | 62.03 | **89.83** | 89.15 | 86.19 | **96.74** | 96.43 | 94.48 |
| Precision | **100.00** | 99.88 | **100.00** | **100.00** | 99.90 | **100.00** | **100.00** | 99.91 | **100.00** |
| Recall | **68.48** | 67.61 | 61.19 | **89.61** | 89.00 | 85.89 | **96.67** | 96.44 | 94.36 |
| Specificity | 99.98 | 96.32 | **100.00** | 99.95 | 95.99 | **100.00** | 99.94 | 96.01 | **100.00** |
| F1-Score | **82.29** | 80.64 | 75.92 | **94.51** | 94.13 | 92.41 | **98.31** | 98.14 | 97.10 |
| MCC | **0.21** | 0.20 | 0.18 | **0.40** | 0.37 | 0.34 | **0.62** | 0.59 | 0.52 |

| Read Accuracy(%) | 95 | | | 98 | | |
|---|---|---|---|---|---|---|
| **Tool** | **Read-Bouncer** | **SPUMONI** | **minimap2** | **Read-Bouncer** | **SPUMONI** | **minimap2** |
| Accuracy | **99.00** | 98.84 | 97.19 | **99.28** | 99.13 | 97.52 |
| Precision | **100.00** | 99.91 | **100.00** | **100.00** | 99.91 | **100.00** |
| Recall | **98.98** | 98.90 | 97.13 | **99.27** | 99.20 | 97.47 |
| Specificity | 99.93 | 95.90 | **100.00** | 99.88 | 95.98 | **100.00** |
| F1-Score | **99.49** | 99.40 | 98.54 | **99.63** | 99.56 | 98.72 |
| MCC | **0.82** | 0.79 | 0.65 | **0.86** | 0.83 | 0.67 |

**Table S5:** Comparing ReadBouncer, SPUMONI, and minimap2 across various metrics on a simulated Zymo Mock Community consisting of seven bacterial species and Saccharomyces cerevisiae. We simulated 360 nucleotide long reads of varying levels of sequence accuracy for all eight organisms. All reads were mapped against the seven bacterial reference sequences to filter out only the bacterial reads. At the same time, we want to keep as much Saccharomyces cerevisiae reads, which corresponds to an enrichment of that organism in a real-world experiment. ReadBouncer can classify a higher percentage of bacterial reads at all levels of read accuracy while having only slightly less specificity than minimap2.
